# Phytoplankton exudates provide full nutrition to a subset of accompanying heterotrophic bacteria via carbon, nitrogen and phosphorus allocation

**DOI:** 10.1101/2021.07.27.453958

**Authors:** Falk Eigemann, Eyal Rahav, Hans-Peter Grossart, Dikla Aharonovich, Daniel Sher, Angela Vogts, Maren Voss

## Abstract

Marine bacteria rely on phytoplankton exudates as carbon sources (DOCp). Yet, it is unclear to what extent phytoplankton exudates also provide nutrients such as phytoplankton-derived N and P (DONp, DOPp). We address these questions by exudate addition experiments from the ubiquitous pico-cyanobacterium *Prochlorococcus* to contrasting ecosystems in the Eastern Mediterranean – a coastal and an open-ocean, oligotrophic station. Nutrient addition did not lower the incorporation of exudate DONp, nor did it reduce alkaline phosphatase activity, suggesting that bacterial communities are able to exclusively cover their nitrogen and phosphorus demands with organic forms provided by phytoplankton exudates. Approximately half of the cells in each ecosystem took up detectable amounts of *Prochlorococcus*-derived C and N, yet based on 16S rRNA sequencing different bacterial genera were responsible for the observed exudate utilization patterns. In the coastal community, several phylotypes of *Aureimarina*, *Psychrosphaera* and *Glaciecola* responded positively to the addition of phytoplankton exudates, whereas phylotypes of *Pseudoalteromonas* increased and dominated the open ocean communities. Together, our results strongly indicate that phytoplankton exudates provide coastal and open-ocean bacterial communities with organic carbon, nitrogen and phosphorus, and that phytoplankton exudate serve a full-fledged meal for specific members of the accompanying bacterial community in the nutrient-poor eastern Mediterranean.

## Introduction

Approximately 50% of the global primary production is executed by oceanic phytoplankton (Field et al., 1998) with > 100 Pg fixed carbon per year (Huang et al., 2021), and generally 50% of the photosynthetically fixed carbon is then consumed by heterotrophic bacteria (Azam and Malfatti, 2007). In addition to utilizing photosynthetically-derived organic carbon, heterotrophic bacteria also require nutrients such as nitrogen and phosphorus (N and P). Some of the N and P is bound to the carbon (C) as part of the dissolved and particulate organic matter released by phytoplankton, and the recycling of these and other micro- and macro-nutrients may directly impact phytoplankton dynamics (Amin et al., 2012; Moore et al., 2013; Buchan et al., 2014; Amin et al., 2015). Thus, oceanic bacteria are key players in biogeochemical cycles with global importance (Azam and Malfatti, 2007; Amin et al., 2015; York, 2018), and interactions between phytoplankton and bacteria may be considered crucial for carbon and nutrient fluxes through aquatic food webs (Cole, 1982; Amin et al., 2015; Christie-Oleza et al., 2017; Seymour et al., 2017; Mühlenbruch et al., 2018). Dissolved organic material (DOM) is released by phytoplankton (DOMp) via passive leakage, active release, as well as through lysis products of dead cells (Grossart, 1999; Thornton, 2014; Christie-Oleza et al., 2017), whereby the type of release may define its composition (Livanou et al., 2017). Besides DOC, heterotrophic bacteria also utilize dissolved organic nitrogen (DON) (Karlson et al., 2015) and phosphorus (DOP) (Riemann et al., 2009) because inorganic forms of both nutrients are scarce in most oceanic environments (Moore et al., 2013; Saito et al., 2014; Liefer et al., 2019). Phytoplankton exudates contain organic nitrogen and phosphorus and serve the accompanying bacterial community as nutrient source (Bertilsson et al., 2005; Roth-Rosenberg et al., 2021a), but their importance to the overall nutrient status of heterotrophic bacteria is still unclear. Also, the role of inorganic nutrients in the utilization of DOCp is not consistent, as inorganic nutrients may (Carlson et al., 2004; Thornton, 2014) or may not (Carlson and Ducklow, 1996) fuel the DOCp utilization by the bacterial community under nutrient limiting conditions, depending on so far unknown factors.

As suppliers of various carbon and nutrient sources, phytoplankton drive bacterial community dynamics (Rooney-Varga et al., 2005), and different phytoplankton release different types of DOMp (Mühlenbruch et al., 2018). Some bacteria are adapted to DOMp derived from specific phytoplankton species (Grossart et al., 2007; Sarmento and Gasol, 2012; Beier et al., 2015) or phytoplankton source communities (Carlson et al., 2004) and thus act as specialists. Others, however, apply generalist strategies and are strongly affected by DOM concentrations (Sarmento et al., 2016). The questions what fraction of the total bacteria are active as well as which exudate compounds are used by them remained open in preceding studies.

In this study, we addressed the following questions: i) Does DOMp serve as a substantial nitrogen and phosphorus source for the accompanying bacterial community? ii) Do bacterial cells selectively incorporate carbon or nitrogen or do they incorporate both at a constant ratio? iii) Which fraction of the total bacterial community is active in DOMp utilization? iv) Which specific bacterial taxa utilize DOMp, and v) How is the bacterial DOCp utilization affected by inorganic nutrients? We explore these questions in two contrasting systems in the Eastern Mediterranean – a coastal and an open ocean station. The pelagic Eastern Mediterranean is ultra-oligotrophic, comparable with major ocean gyres (Hazan et al., 2018; Reich et al., 2021), whereas conditions closer to the coast are somewhat richer in nutrients (Sisma-Ventura and Rahav, 2019). As a DOM source, we used spent media from Prochlorococcus strain MIT9312, labelled with ^13^C and ^15^N. The cyanobacterium *Prochlorococcus* is the most abundant phototrophic organism on Earth, with an annual global mean abundances of 2.9 x 10^27^ cells (Flombaum et al., 2013), and dominates phytoplankton biomass in many oligotrophic oceans despite its small size (Partensky et al., 1999). *Prochlorococcus* alone was suggested to exude as much as 75% of the daily photosynthetic organic carbon production in oligotrophic environments (Ribalet et al., 2015), resulting in up to 40% of the total bacterial production (Bertilsson et al., 2005; Biller et al., 2015). In the Eastern Mediterranean, the structure of the free-living heterotrophic bacterial community is weakly but significantly correlated with the presence of divinyl Chlorophyll A, a diagnostic pigment of Prochlorococcus, further supporting a potential link through DOM production and uptake (Roth-Rosenberg et al., 2021b). The specific strain used, MIT9312, was selected because it is the most abundant ecotype globally (Johnson et al., 2006), although not in the Mediterranean (Mella-Flores et al., 2011), and has recently been show to exude large amounts of DOC (Roth-Rosenberg et al., 2021a). To test for the above raised questions, we amended coastal and open-ocean bacterial communities with MIT9312 exudates with and without additions of inorganic nutrients, and analysed bacterial responses via 16S rRNA amplicon sequencing, cell numbers, bacterial production, incorporation of DOCp and DONp, and alkaline phosphatase activity.

## Results

### Contrasting conditions at the coastal vs. open-ocean sites

The two sampling sites exhibited distinctively different chemical characteristics. The inorganic nutrient concentrations at the coastal site under influence of the storm event were 0.12 ± 0.01 µM PO_4_^3-^, 3.00 ± 0.63 µM NO_2+3_^-^, and 0.50 ± 0.11 µM NH_4_^+^. In contrast, the open ocean site was highly oligotrophic even though the samples were taken during winter when the water column was relatively well mixed; <0.006 µM PO_4_^3-^ and 0.35 µM NO_2+3_^-^ (Reich et al., 2021).

### Cell numbers and cell-specific bacterial production

At the start of the experiment (t0), the coastal bacterial community showed approximately 5-times higher cell numbers compared to the open-ocean community (Fig. 1a and b). After 24 h, cell numbers significantly decreased in the control + nutrients and exudate treatments in the coastal community (Fig. 1a), whereas in the open-ocean community, both exudate treatments (with and without nutrients) showed elevated cell numbers (∼ 1.5-fold), which were, however, not significant (Fig. 1b). Thus, neither the nutrient nor the exudate additions resulted in a clear impact on cell numbers after 24 h of incubation.

**Fig. 1:**
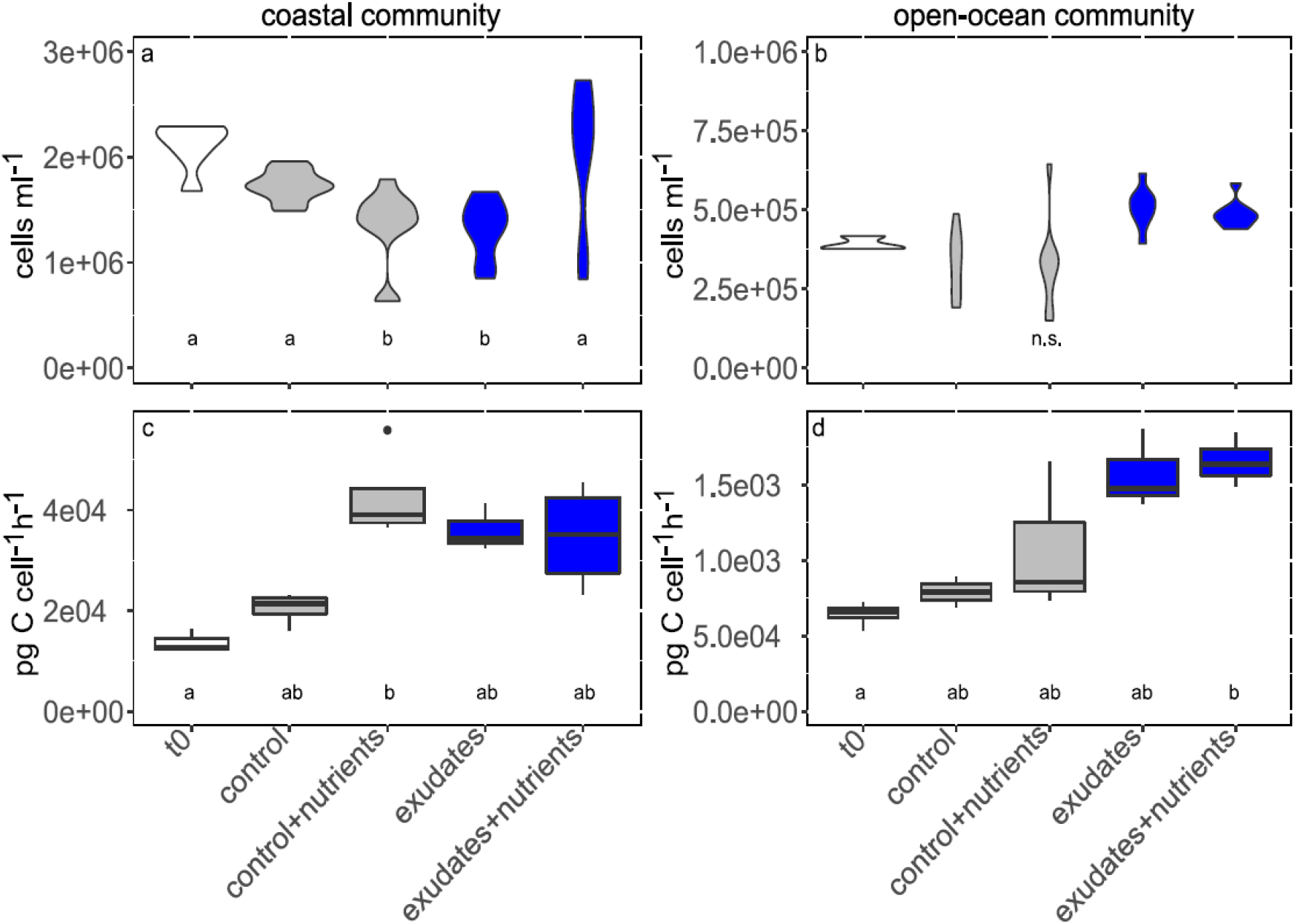
Cell numbers (violin plots, top row) and bacterial production (box-plots, bottom row) in the different treatment microcosm bottles. The letters in the panels represent the outcomes of Tukey post-hoc tests. Please note the different y-scales for the coastal and open-ocean communities. T0 samples are displayed in white, control samples in grey, and exudate samples in blue.

Cell-specific bacterial production was more than 4-times higher in the open-ocean community when compared to the coastal community (Fig. 1c and d), whereas bulk measurements revealed similar outcomes in both environments (Supplemental 1). Although the cell-specific bacterial production increased in all treatments after 24 h, it differed significantly from the control only in the nutrient amendment for the coastal community (Fig. 1c) and in the exudate + nutrient treatment in the open-ocean community (Fig. 1d). The increase in cell-specific bacterial production in the control + nutrient and exudate treatments may partly be caused by increased cell sizes (FSC-A as cell size proxy) for the coastal community, but this did not apply for the open-ocean community (Supplemental 2).

### Incorporation of organic carbon and nitrogen

In order to verify that bacteria incorporate carbon and nitrogen derived from DOMp, we investigated their uptake by means of NanoSIMS measurements. Both coastal and open-ocean bacterial communities showed significant incorporation of labelled organic carbon and nitrogen derived from the phytoplankton exudates. As expected, the coastal community at time 0 and all control treatments revealed values around the naturally occurring ratios of 0.011 (^13^C/^12^C) and 0.00367 (^15^N/^14^N)(Fig. 2, no t=0 samples were available for the open-ocean location). Nutrient additions had neither an effect on the DOCp (Fig. 2a and b), nor on the DONp incorporation (Fig. 2c and d). After 24 h of incubation, in the coastal bacterial community 54% and 47% of the cells in the exudate and exudate + nutrients treatments revealed increased ^15^N and ^13^C values, respectively, whereas in the open-ocean community 39% and 51% cells with increased ^15^N and ^13^C values were found (increased ratios defined as ratios above the 95% percentile of pooled t0, control and control + nutrient measurements, Supplemental 3). In the coastal as well as in the open-ocean bacterial community ^13^C and ^15^N uptake were highly correlated (Table 1). If the single treatments were considered separately, for both, the coastal and open-ocean communities both exudate treatments revealed significant correlations, but as expected, none of the other treatments (Table 1). However, the slopes of the exudate treatments were steeper in the treatments with on-top additions of nutrients (Table 1).

**Fig. 2:**
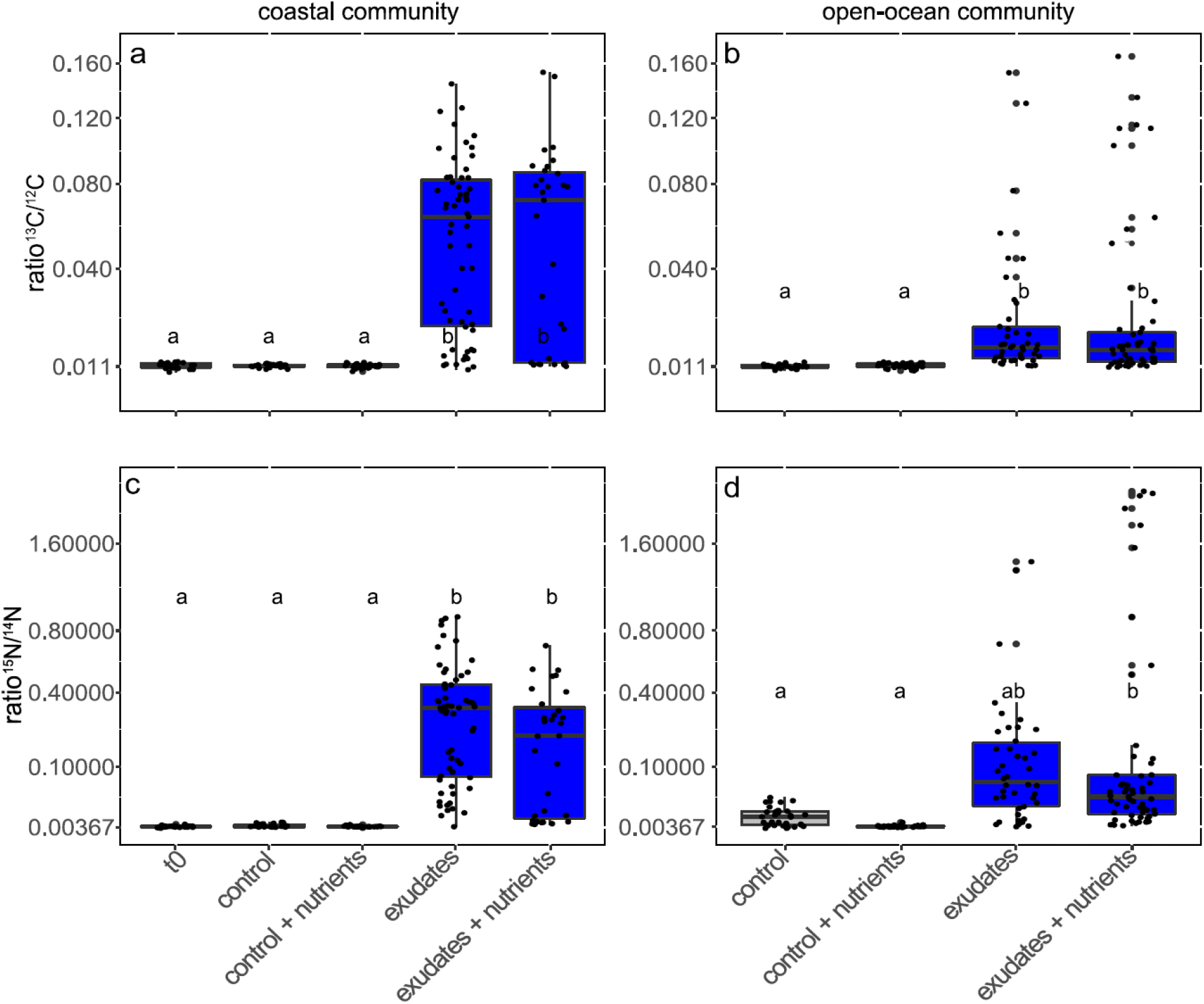
^13^C/^12^C ratios (A, B) and ^15^N/^14^N ratios (C, D) of the coastal (A, C) and the open-ocean (B, D) bacterial community following 24 hrs incubations and in t0. T0 samples are displayed in white, control samples in grey, and exudate samples in blue.

**Table 1:** Correlations between ^13^C and ^15^N incorporation derived from NanoSIMS analyses. The y-axis was defined as = ^13^C/^12^C ratio, the x-axis as = ^15^N/^14^N ratio.

| Experiment | Treatment | Linear function | r<sup>2</sup> | p |
| --- | --- | --- | --- | --- |
| Exp. 1, coastal | All treatments | - | 0.53 | < 0.0001 |
|  | t0 | - | -0.03 | 0.63 |
|  | control | - | 0.01 | 0.39 |
|  | control + nutrients | - | -0.1 | 0.05 |
| | exudates | $y = 0.23x$ | 0.72, | <0.0001 |
| | exudates + nutrients | $y = 0.20x$ | 0.48 | <0.0001 |
| Exp. 2, open-ocean | All treatments | - | 0.91 | <0.0001 |
|  | control | - | -0.03 | 0.77 |
|  | control + nutrients | - | -0.02 | 0.78 |
| | exudates | $y = 0.24x$ | 0.94 | <0.0001 |
| | exudates + nutrients | $y = 0.15x$ | 0.95 | <0.0001 |

### Cell-specific alkaline phosphatase activity

To test whether bacterial communities satisfy their phosphorus demand via DOMp, we analysed the activity of the alkaline phosphatase enzyme (APA) in the different treatments and environments. At time 0, cell-specific APA was approximately seven times higher in the open-ocean community compared to the coastal community (Fig. 3), whereas bulk (i.e. per volume seawater) analyses showed comparable outcomes in both environments (Supplemental 5). In both environments, additions of phytoplankton exudates lowered the alkaline phosphatase activity, without any effect of on top additions of nutrients. Nutrient additions in the controls also lowered the APA, but not as strongly as the exudate additions (Fig. 3). In the coastal bacterial community, the control treatments without nutrient additions revealed a significantly higher cell-specific APA compared to all other treatments, and in the open-ocean community the exudate treatments had not only lower APA compared to the control treatment, but also lower APA compared to the t0 and control + nutrient treatments (Fig. 3).

**Fig. 3:**
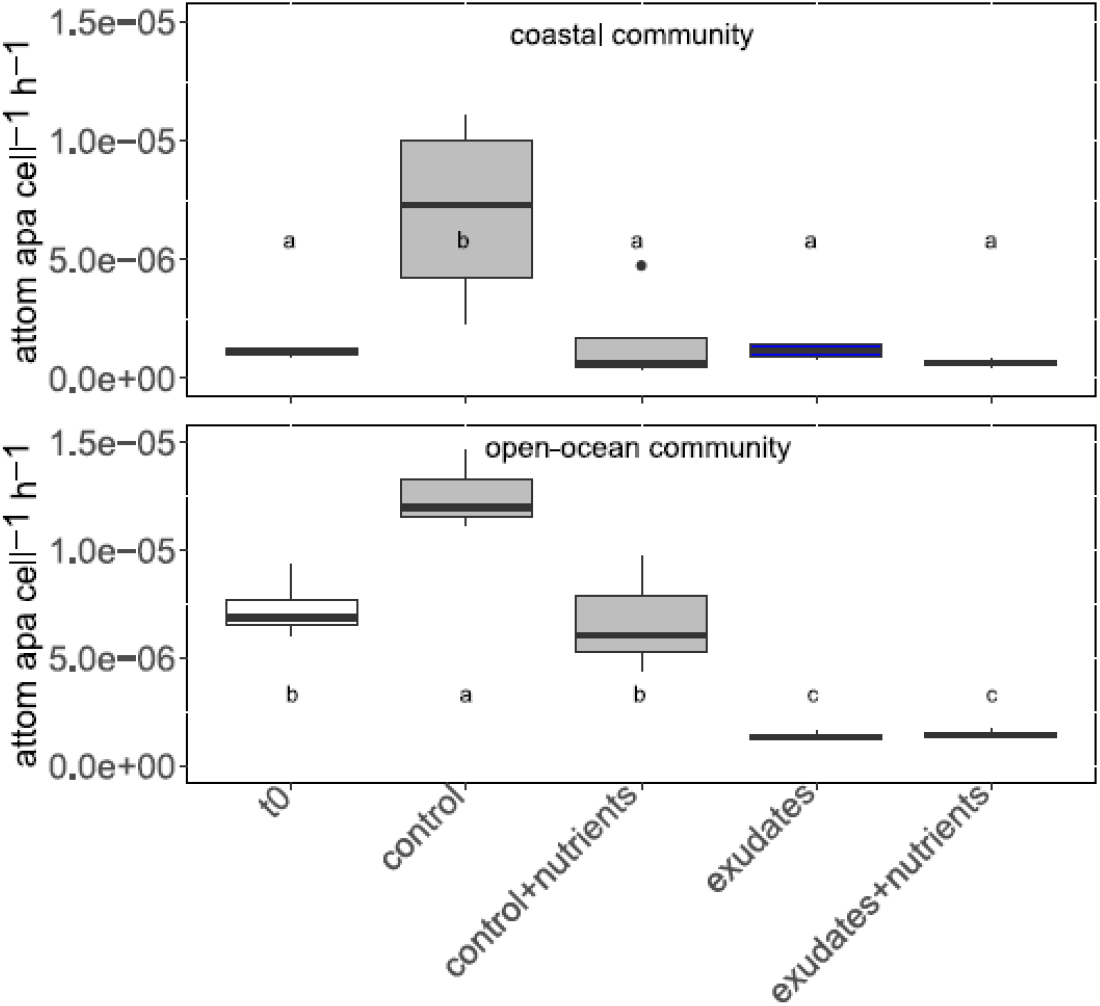
Cell-specific alkaline phosphatase activity for the coastal (top panel) and the open-ocean (bottom panel) bacterial community. The letters in the panels represent the outcomes of Tukey post-hoc tests. T0 samples are displayed in white, control samples in grey, and exudate samples in blue.

### Dynamics in bacterial community composition

To answer how the active bacterial communities develop in response to exudate additions, we analysed the diversity and composition in the 16S rRNA derived dataset. Since the amplicons were derived from RNA molecules, these represent primarily the active members of the bacterial community. Shannon diversities did not show significant differences between the treatments in the coastal communities (ANOVA, p = 0.81), but in the open-ocean community exudate treatments yielded lower Shannon diversities (ANOVA, p = 0.00, Fig. 4).

**Fig. 4:**
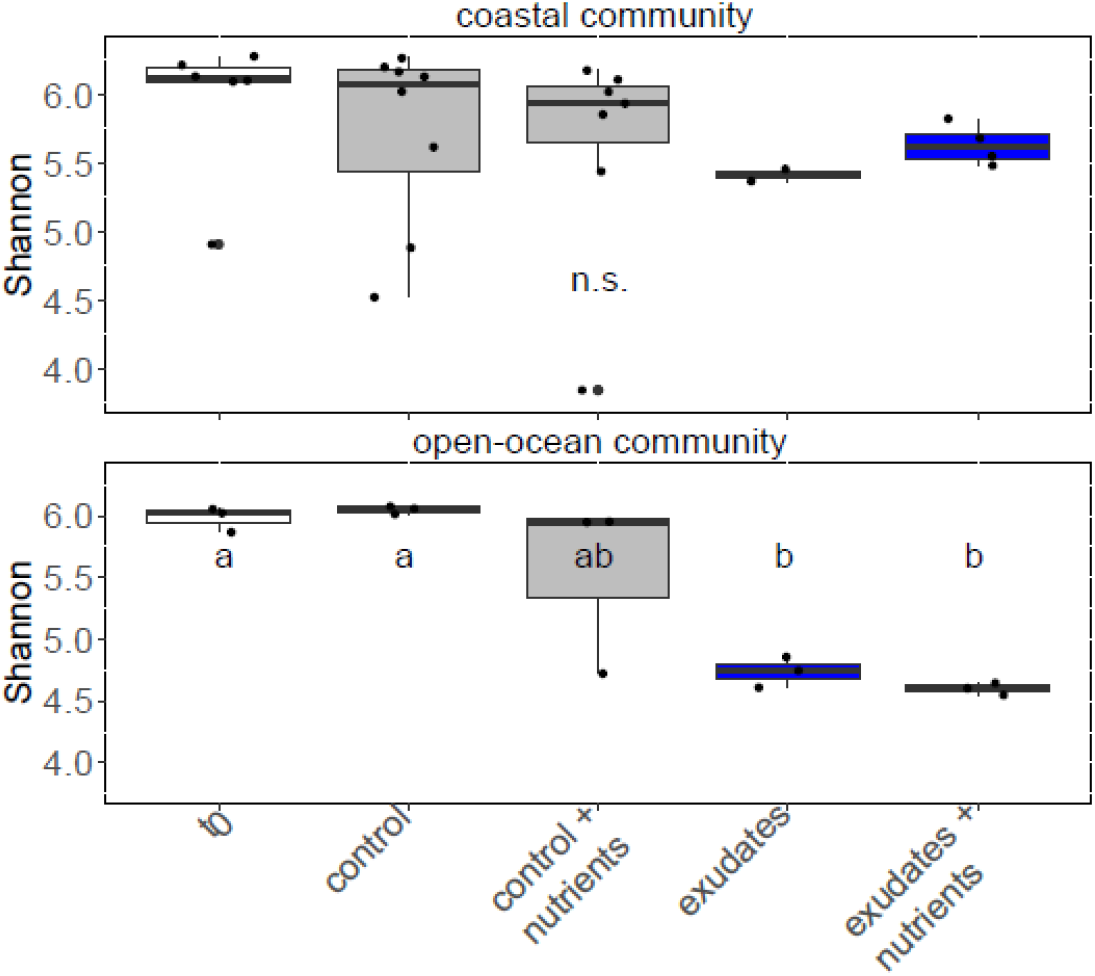
Shannon diversity for the coastal (top panel) and the open-ocean (bottom panel) bacterial community. The letters in the panels represent the outcomes of Tukey post-hoc tests. Please note that in coastal communities size-fractionated (5 and 0.2 µm pore-width) samples were analysed, whereas in the open-ocean communities no size-fractionation was performed (only 0.2 µm filter pore-width). T0 samples are displayed in white, control samples in grey, and exudate samples in blue.

NMDS analyses of the bacterial communities showed differences between experiments (ANOSIM, p=0.001, R=0.44, Fig. 5), and in the coastal community additionally between the free and attached fractions (ANOSIM, p=0.001, R=0.45, Fig. 5). However, the different treatments also clustered significantly differently (ANOSIM, p=0.02, R=0.11), where strong differences occurred between samples with added exudates and those without. The addition of nutrients did not show any clear effects in either experiment (Fig. 5). If only the 40 most abundant ASVs in both experiments were considered and illustrated in heatmaps, likewise strong community shifts were observed between the treatments (Fig. 6a and b). LEfSe (linear discriminant analysis effect size) analyses suggested *Fluviicola*, *Pseudofulvibacter*, *Synechococcus* and NS4 marine group being predominant in t0 samples, *Spongiispira* predominant in the controls, whereas the exudate treatments were dominated by *Glaciecola* in the coastal environment (Supplemental 5, Fig. 6a, see also the 20 most abundant ASVs for each single treatment (Supplemental 6)). In the open-ocean community, *Pseudoalteromonas* was strikingly abundant in the exudate treatments, whereas SAR11, SAR202 and KI89a clade bacteria were predominant at t0 and in the control samples (Fig. 6b, Supplemental 6, Supplemental 7).

**Fig. 5:**
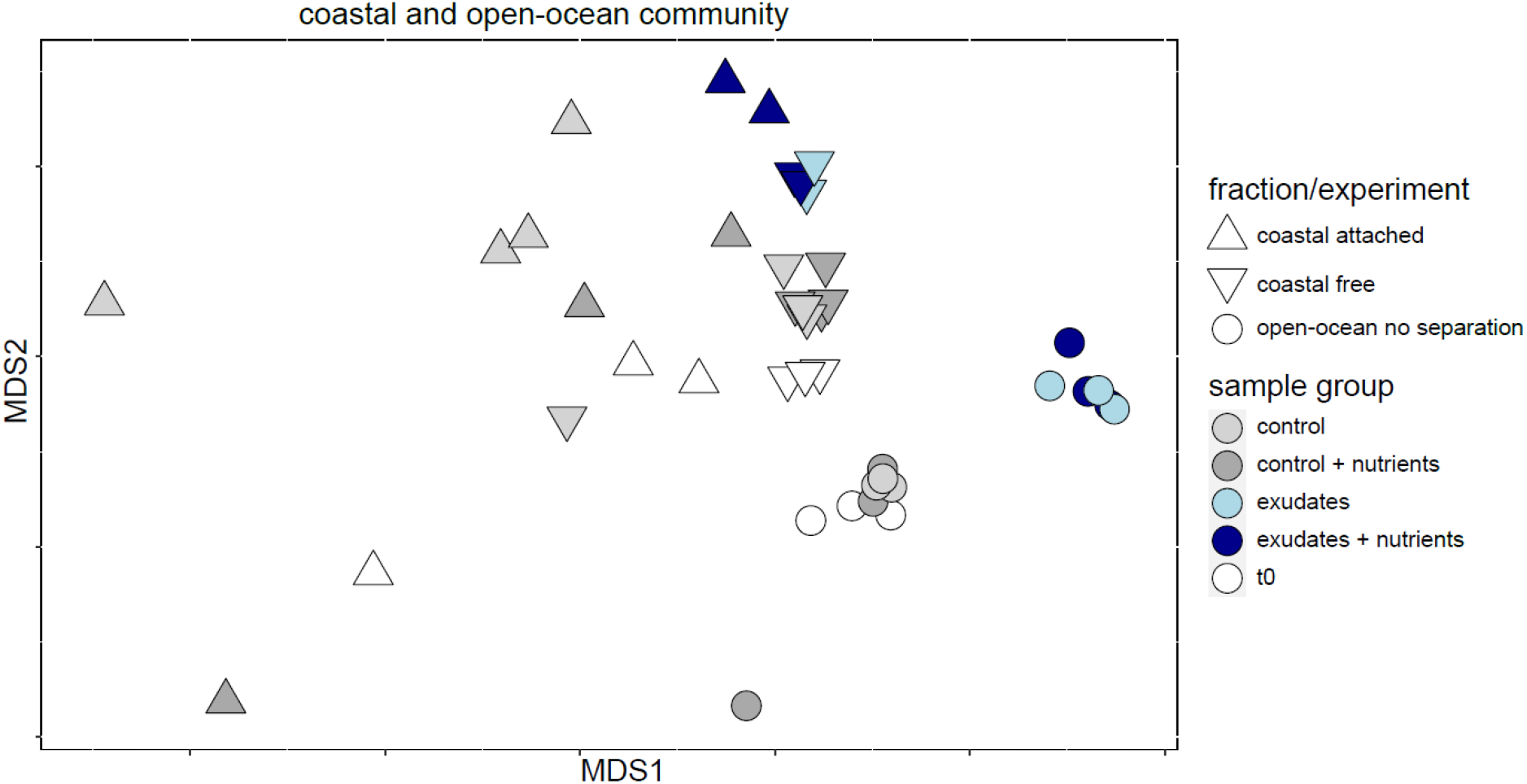
NMDS plot of the bacterial communities (stress = 0.15). The symbol and colour key is given on the right-hand side. T0 samples are displayed in white, control samples in grey, and exudate samples in blue.

**Fig. 6:**
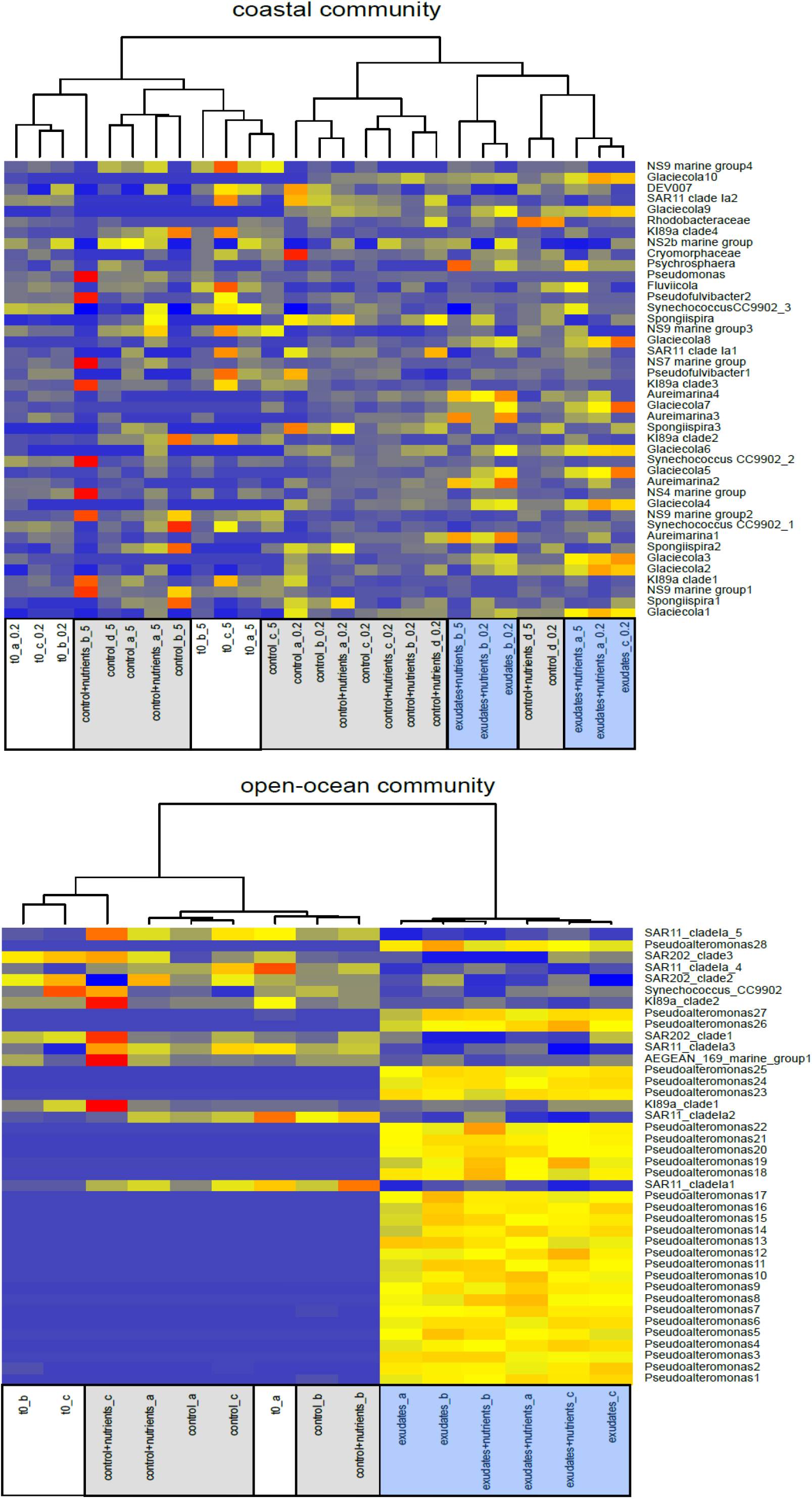
Heat-maps of the 40 most abundant ASVs (normalized absolute read counts) in the coastal (a) and open-ocean (b) environment. Samples can be seen below the heatmap, ASVs on the right side. If several ASVs with the same taxonomy were present, the taxonomy was numbered. Blue colour indicates low read numbers, red colour indicates high read numbers. The letter behind the treatment names refer to the replicate, the number behind this letter indicates the filter pore-width and thus the fraction (only coastal experiment, 0.2 µm = free fraction, 5 µm = attached fraction). Dendrograms on top are calculated based on Euclidian distances, the ordering of ASVs refer to read counts (top ASV: lowest read counts, bottom ASV: highest read counts). T0 samples are displayed in white, control samples in grey, and exudate samples in blue.

## Discussion

In this study, we wanted to identify which fraction (and which bacterial taxa) of the total bacterial community are active in DOMp utilization, whether DOMp serves as a substantial nitrogen and phosphorus source for the accompanying bacterial community, and if so, whether bacterial cells selectively incorporate carbon or nitrogen. Our experimental results strongly indicate that DOMp serves as substantial N and P source for a subset of approximately 50% of the bacterial community at the times and places sampled, and that the active community members can fully satisfy their demands with the phytoplankton derived organic nutrient sources. This stands in contrast to other studies from this region, suggesting that the Eastern Mediterranean Sea bacterial communities are limited by inorganic nutrients, predominantly PO_4_ (Krom et al., 2010; Krom et al., 2014). Treatment-specific correlations between carbon and nitrogen incorporation also suggest that both compounds could be selectively incorporated. Another raised question was which factors limit heterotrophic production: Are marine bacteria predominantly carbon limited (Christie-Oleza et al., 2017), or is there ample DOCp for them but not enough nutrients for its utilization (Carlson et al., 2004), because DOMp degradation by bacteria and bacterial production is affected by nutrient availability (Fouilland et al., 2014)? For this point, our results indicate that in our experimental set-ups the necessitated nutrients for the DOCp utilization by the bacterial community were allocated with organic forms included in the DOMp.

### Bacterial production and carbon incorporation following phytoplankton exudate addition

Increased bacterial production in both environments following exudate additions (Fig. 1) suggest that exudate DOCp provide carbon for both bacterial communities, which was confirmed by NanoSIMS measurements (Fig. 2). However, as stated above, DOCp incorporation might be limited by inorganic nutrients (Carlson et al., 2004; Fouilland et al., 2014). For example, the conversion of carbon from polysaccharides into bacterial biomass was enhanced by inorganic nitrogen for *de novo* synthesis of cellular proteins (Grossart et al., 2007; Piontek et al., 2011), and additions of inorganic nitrogen and high nitrate concentrations increased glucose assimilation by bacterioplankton (Bianchi et al., 1998; Skoog et al., 2002). Our results, however, suggest that the nutrient demand for DOCp utilization in both bacterial communities was satisfied with organic sources derived from *Prochlorococcus* exudates because nutrient additions on top of the exudates did not result in higher cell-specific bacterial productivity nor in higher incorporation rates of ^13^C-labelled carbon (Figs. 1 and 2, also see the next paragraph). We observed lower cell-specific bacterial production, but higher ^13^C/^12^C ratios in the coastal community if compared to the open-ocean community (Figs. 1 and 2). The lower cell-specific bacterial production in the coastal community might partly be explained with the storm event that introduced sediment and soil bacteria to the water column. The resuspension might have contributed significantly to our cell counts but not the cell specific activity, lowering only the cell-specific bacterial production. This assumption is supported by (Raveh et al., 2015), where much lower bacterial counts (∼5×10^5^ cells ml^−1^ in January) but comparable bacterial bulk productions were determined for a coastal area in the Eastern Mediterranean Sea. The higher ^13^C/^12^C ratios in the coastal communities, on the other hand, appear surprising taking into account the storm event with possible input of carbon and contradict previous studies, where direct carbon coupling between phytoplankton and bacteria was weak in coastal waters due to substantial allochthonous carbon sources (Morán et al., 2002). Yet, our findings suggest an preferential utilization of DOMp compared to other carbon sources, which fits with an observation of coastal bacterial communities in the Mediterranean satisfying more than half of their carbon demand from phytoplankton exudates (Fouilland et al., 2014). This can be explained by the fact that pico-cyanobacteria (i.e. *Prochlorococcus* and *Synechococcus*) exudates contain high fractions of low molecular weight DOC, which is highly labile and can be rapidly utilized by heterotrophic bacteria (Bertilsson et al., 2005; Sharma et al., 2014), e.g. organic acids, organo-halogens, and isoprene (Shaw et al., 2003; Bertilsson et al., 2005) as well as lipids, proteins, and fragments of DNA and RNA (Biller et al., 2015). The strain *Prochlorococcus* MIT9312 used in our experiments, releases ∼ 90% of the fixed carbon as DOCp (Roth-Rosenberg et al., 2021a).

### Incorporation of DONp and DOPp

Our NanoSIMS measurements as well as APA analyses suggest that the coastal and open-ocean bacterial communities cover their nitrogen and phosphorus demands via DOMp (Figs. 2 and 3). This notion is in accordance with previous studies, showing that pico-cyanobacteria produce nitrogen and phosphorus-rich DOMp (Beliaev et al., 2014; Christie-Oleza et al., 2017) that serves as energy and also nutrient source for bacteria (Livanou et al., 2017). For example, bacteria exposed to phytoplankton exudates upregulated transcription of nitrogen and phosphorus utilization genes (McCarren et al. 2010), and co-cultures of picocyanobacteria with several different heterotrophic bacteria revealed an increased expression of genes for transporters of amino acids and peptides (Beliaev et al., 2014). It has been previously shown that *Prochlorococcus* MIT9312 exudes considerable amounts of organic nitrogen under laboratory conditions (Roth-Rosenberg et al., 2021a), possibly in the form of proteins, amino acids, DNA, RNA and nucleotides. This could provide labile DONp to the bacterial community (Sharma et al., 2014). In general, phytoplankton exudates provide manifold organic nitrogen species to heterotrophic bacteria, including urea, dissolved and free amino acids, proteins, nucleic acids, amino sugars (Meon and Kirchman, 2001; Berman and Bronk, 2003), and methylated amines (Lidbury et al., 2015), etc. In co-culture and exudate addition experiments, especially dissolved free amino acids and urea were used by bacteria (Grossart and Simon, 2007; Bradley et al., 2010; Sarmento et al., 2013; Beliaev et al., 2014). Similar to nitrogen, phytoplankton exudates may offer substantial amounts of organic phosphorus in various forms (Livanou et al., 2017), and our results indicate that both bacterial communities appease their phosphorus demand with the provided phytoplankton exudates (Fig. 3). A previous modeling study has suggested that, under conditions of N starvation, *Prochlorococcus* releases P-containing molecules such as nucleobases and nucleosides (Ofaim et al., 2021), although the magnitude of exudation of DOP is not well constrained (Roth-Rosenberg et al., 2021a). Increased APA in the control incubations indicate phosphorus depletion in these treatments, which is consistent with the low initial phosphorus concentrations and our “batch-like” bottle incubations. Indeed, we could show in the open-ocean community a significantly lowered APA after exudate additions compared to the control + nutrient treatments, indicating not only a complete phosphorus supply by the exudates but a preferential utilization (Fig. 3). In summary, our results strongly suggest that phytoplankton derived organic nitrogen as well as phosphorus is incorporated into bacterial biomass and provide the primary nitrogen and phosphorus source for the coastal as well as open-ocean bacterial communities. Our experimental conditions, however, do not fully mimic natural conditions, for example, the added organic matter is at relatively high concentration, and comes from a single course (a single phytoplankton strain). Further research is required in order to determine whether these patterns also occur when considering naturally-derived phytoplankton exudates in natural environments.

### Relationships between ^13^C and ^15^N incorporation

Ratios of ^13^C to ^15^N incorporation suggest a volitional and selective uptake of organic nitrogen by both communities if inorganic nitrogen is scarce (exudate + nutrient treatments revealed steeper slopes compared to the exudate only treatments with ^15^N = x-axis, ^13^C = y-axis). On the other hand, the highest ^13^C to ^15^N uptake ratios occurred in the open-ocean community in the exudate + nutrient treatments, indicating a carbon limitation and/or luxury uptake of nitrogen in this community. This stands in stark contrast with suggestions that especially open-ocean microorganisms minimize their nitrogen costs due to nitrogen limitation (Grzymski and Dussaq, 2012). However, bacterial communities show distinct seasonal changes in nutrient limitation, and our experiments illustrate a single time-point of a “winter”-community, in which carbon might be the predominate limiting factor (Pinhassi et al., 2006). Therefore, we cannot rule out co-limitation of carbon and nitrogen, which, in fact may have occurred (Rahav et al., 2019). Altogether, the ratios of ^13^C to ^15^N incorporations derived from NanoSIMS analyses in our experiments were in the same range as for phytoplankton associated bacteria in the Baltic Sea (Eigemann et al., 2019), and different ^13^C to ^15^N ratios in NanoSIMS measurements of different treatments suggest selective uptakes of either carbon or nitrogen.

### Community responses to exudate additions

In the open-ocean communities, as a reaction to exudate additions a decrease in Shannon diversity was obvious (Fig. 4), which was caused by the dominance of a single genus, namely *Pseudoalteromona*s (Fig. 6, Supplemental 6), whereas diversity in the coastal community remained constant, but with different abundant genera in the different treatments (Figs. 4 and 6). However, one should keep in mind that our analyses are based on rRNA, and thus only the active part of the bacterial community is appropriately reflected. Together, after exudate additions, *Pseudoalteromonas* in the open-ocean communities, and *Glaciecola, Psychrosphaera* and *Aureimarina* in the coastal communities revealed strong positive responses (Fig. 6, Supplemental 5, 6, 7). *Pseudoalteromonas* belongs to the order Alteromonadales which significantly contribute to carbon cycling in the surface ocean (Pedler et al., 2014), possesses effective degradation systems for a wide array of polysaccharides (Gobet et al., 2018), and showed positive responses to phytoplankton exudates (Seymour et al., 2009). *Glaciecola* belongs to the Alteromonadaceae, which were likewise enriched upon phytoplankton exudate addition (Taylor and Cunliffe, 2017). In general, the responsive bacteria in our experiments possess a variety of carbohydrate-active enzymes (CAZymes). For example, Glaciecola sp. 4H-3-7YE-5 possesses the genomic possibility for the degradation of several polysaccharides (Klippel et al., 2011), which constitute major components of phytoplankton exudates (Meon and Kirchman, 2001; Mühlenbruch et al., 2018). Further, cyanobacteria are known to produce glycogen as a storage polysaccharide (Bertocchi et al., 1990; Bhatnagar and Bhatnagar, 2019), and *Pseudoalteromonas*, *Psychrosphaera* as well as *Glaciecola* possess effective utilization systems for glycogen (Lombard et al., 2014), and other polysaccharides common in phytoplankton (Klippel et al., 2011; Pheng et al., 2017; Gobet et al., 2018).

Despite that bacterial communities were comparatively similar at t0 in coastal and open-ocean environments, different responders to exudate additions appeared (Fig. 5, Supplemental 5, 6, 7). Below, we list four possible explanations for the different development after exudate additions: Firstly, the source communities differed altough several abundant members overlapped. Indeed, *Pseudoalteromonas*, the main responder in the open-ocean treatments, was even more relatively abundant at t0 in the coastal community (mean subsampled read-sums of *Pseudoalteromonas* ASVs in t0 samples: 5 for open-ocean and 19 for coastal communities, Supplemental 8), but the most responsive genera of the coastal community were completely lacking in the open-ocean community (*Psychrosphaera, Aureimarina*, Supplemental 8) or present at low abundances (*Glaciecola*, Supplemental 8). This outcome suggests a high importance of the bacterial source communities on the effectiveness in DOC utilization as well as the ability to use different DOC sources (Carlson et al., 2004; Grossart et al., 2007). Secondly, our taxonomic resolution did not go beyond the genus level, and as consequence open-ocean and coastal communities may have been indeed more dissimilar than suggested by our analyses at t0 (i.e. different species and strains of the same genus between open-ocean and coastal bacterial communities), ultimately resulting in an underestimation of differences between source communities between environments. Thirdly, environmental conditions favour certain bacterial genera over others (Nemergut et al., 2013; Wu et al., 2019), which may reflect the ability to effectively utilize DOMp. Thus, under open-ocean conditions paired with the additions of DOMp, *Pseudoalteromonas* was the most successful genus that outcompeted other genera, whereas under coastal conditions it was outcompeted, despite its higher relative abundances at t0. Our data suggest that other members in the open-ocean community were not able to show such a strong response in the 24 h of incubation, and Alteromonadales are known as opportunists (Eilers et al., 2000) and fast growers (Pedler et al., 2014). Fourthly, the dominant response of the copiotrophic generalist *Pseudoalteromonas* (Gammaproteobacteria) in the open-ocean environments may be explained with drastic changes in relative DOM concentrations, whereas in the coastal environment additions of DOM induced more specialists responses (Sarmento et al., 2016). Absolute DOC concentrations are an important factor for the uptake ability of heterotrophic bacteria, as only a few specialists were able to incorporate DOCp at low concentrations, but a broader range of bacteria could use the same source of DOCp at high concentrations (Sarmento et al., 2016). This might be also true in our experiments, where the background DOC concentrations probably have been much higher in the coastal (especially after the storm event) compared to the open-ocean environment.

Besides specific outcomes at the genus level, some general pattern occur under phytoplankton bloom conditions, with Flavobacteria, Alphaproteobacteria and Gammaproteobacteria being dominant classes (Buchan et al., 2014). This is partly reflected in our experiments (Fig. 6), with especially Gammaproteobacteria (*Pseudoalteromonas, Psychrosphaera, Glaciecola*), and Flavobacteria (*Aureimarina*) showing strong positive responses to exudate additions, whereas we did not observe positive responses but rather relative declines of Alphaproteobacteria (Supplemental 6). This relative decline of Alphaproteobacteria as response to phytoplankton exudates partly contradicts previous studies, where especially the Roseobacter clade (class Alphaproteobacteria) reveals numerous positive interactions with phytoplankton (Romera-Castillo et al., 2011; Lidbury et al., 2015). However, despite being capable of using a wide array of substrates (Lidbury et al., 2015), Roseobacter did not take up *Prochlorococcus* exudates in a similar experiment (Sarmento and Gasol, 2012), emphasizing the selective uptake of DOMp by different bacteria (Sarmento et al., 2013).

A point not addressed in our experimental set-up is the successive degradation from labile to recalcitrant DOM. In general, DOMp utilization by heterotrophic bacteria undergoes a succession with different responsive organisms at different times after DOMp pulses (Teeling et al., 2012; Buchan et al., 2014; Teeling et al., 2016). Oligotrophic organisms like SAR11 utilize highly labile low molecular weight compounds, whereas copiotrophic bacteria such as Alteromonadales utilize high molecular weight compounds such as polysaccharides (Sharma et al., 2014). With our experiments, we only reflect the community response at a single time-point, i.e. 24 h after DOMp addition, which may resemble daily changes in DOMp concentrations accompanied with photosynthesis during phytoplankton blooms.

Furthermore, as mentioned above, *Prochlorococcus* exudes high amounts of labile DOMp, which can be rapidly utilized by heterotrophic bacteria (Roth-Rosenberg et al., 2021a). Thus, the concentration of the added DOC as well as the incubation time allow for a maximal comprehensive picture (for a single pulse), and thus should reflect responses of oligotrophic as well as copiotrophic community members at environmental relevant concentrations (McCarren et al., 2010; Seymour et al., 2010; Sarmento and Gasol, 2012; Sharma et al., 2014; Beier et al., 2015; Sarmento et al., 2016). However, to fully comprehend the dynamics of DOCp utilization and the community succession additional time-course studies are needed.

## Conclusions

Short-term responses of coastal and open-ocean bacterial communities to phytoplankton exudates addition with and without inorganic nutrients revealed similar bacterial response patterns, but different responders in the coastal vs the open ocean communities. The different responders suggest environmental factors, such as the ambient DOM concentrations or the initial bacterial communities to determine DOMp utilization effectiveness of the respective bacterial communities to some extent. Our results strongly indicate that the allocated phytoplankton exudates provide a major fraction of the bacterial community (∼50%) with organic carbon, nitrogen and phosphorus as combined additions of exudates and inorganic nutrients neither enhance cell-specific bacterial production, nor lowered incorporation of DONp and cell-specific alkaline phosphatase activity. Utilization of DOCp seems not to be limited by inorganic nutrients, because addition of inorganic nutrients did not elevate the incorporation of DOCp into bacterial cells. Consequently, phytoplankton exudates may function as a full-fledged meal for the accompanying bacterial communities, and can be used as energy and nutrient sources by bacteria independent of the surrounding inorganic nutrient concentrations. *Prochlorococcus* is a major component of the global phytoplankton (Flombaum et al., 2013), and its exudates likely substantially contribute to bacterial production in oligotrophic environments (Biller et al., 2015; Ribalet et al., 2015). Hence, together with the naturally occurring amount of added DOM (in bloom occasions) (Sharma et al., 2014; Beier et al., 2015), our results may reflect important patterns in marine environments. Together, our study emphasizes the dependency of heterotrophic bacteria on phytoplankton exudates, and illustrates that abiotic factors may resign beyond biotic interactions for marine heterotrophic bacteria. Therewith, our outcomes further strengthen the importance of phytoplankton-bacteria interactions in carbon, nutrient and mineral cycling, and thus in functioning of marine ecosystems.

## Experimental procedures

### Experimental set-up

*Prochlorococcus* MIT9312 was grown under constant light (20 μ E m^−2^ sec^−1^) at 22°C in Pro99 media where the NH_4_ concentration was reduced from 800 µM to 100 µM, resulting in the cells entering stationary stage due to N starvation (Grossowicz et al., 2017). For several generations before harvest, 98% labelled ^15^N-NH_4_^+^ was used as the sole N source, and the media was amended with 1 mM of 98% labelled ^13^C-HCO_3_ as a C source, resulting in a fully-labeled culture. To obtain cell-free exudates, batch cultures were grown as described above in 2L bottles, and were harvested at the early decline phase by centrifugation followed by filtration through a 0.22 µm polycarbonate filter. Early stationary phase was chosen to minimize the carryover of ^15^N-NH_4_^+^ (which should be depleted from the media, see below), to increase the amount of released DOC (Roth-Rosenberg et al., 2021a), and because spent media from this stage may reflect natural DOMp composition better than DOMp from exponentially growing cultures (Christie-Oleza et al., 2017). We note that the DOM in the media is likely a result of both exudation and cell mortality. The spent media was maintained at −20°C until use.

Using the ^15^N and ^13^C labelled spent media we performed two incubation experiments in order to test the bacterial response to DOMp in coastal (Exp 1) compared to open-ocean (Exp 2) systems. Each of these experiments included the spent media with and without inorganic nutrient additions of 20 µM NH_4_, and 2 µM PO_4_, to eliminate nutrient limitation in the 24 h of exposure in the nutrient spiked treatments (Table 2). Exp 1 was carried out in the dark on Jan. 9^th^, 2019 in 1 m³ natural seawater flow-through tanks to maintain ambient temperature at the Israel Oceanographic and Limnological Research centre in Haifa, Israel. Each treatment consisted of 4 biological replicates. The coastal site was a 5 m intake pipe during a winter storm with high waves and significant turbulence. Additionally, heavy rainfalls caused considerable land-to-sea run-offs. As a result, the water was brownish in colour, which may have added allochthonous material, as well as soil and sediment bacteria. Exp 2 was carried out on board of the R/V Mediterranean Explorer on Jan. 23^rd^, 2019 at station THEMO-2 (Reich et al., 2021), in an on-board flow-through system, and each treatment consisted of 3 biological replicates. For both experiments, 4.5 l Nalgene bottles were filled with 3 l of 50 µm pre-filtered sea water with the respective DOCp and nutrient amendments (Table 2). In order to receive a clear response of the bacterial community in the range of naturally occurring concentrations of DOC, we chose the addition of 25 µM DOCp (Seymour et al., 2010; Beier et al., 2015; Sarmento et al., 2016). Likewise, the incubation time was set to 24 h, in order to see a meaningful response of both, fast and slow responsive bacteria to phytoplankton exudates at the above-mentioned DOC concentration (McCarren et al., 2010; Sarmento and Gasol, 2012; Sharma et al., 2014; Beier et al., 2015). The spent *Prochlorococcus* medium contained 0.9 µM PO_4_ and 0.16 µM NH_4_, resulting in a final concentration of 5 nM ^15^NH_4_^+^ in the exudate treatments (32x diluted). We accounted for these labelled, inorganic leftovers in the control and control + nutrient treatments in order to differentiate between uptake of DOMp and inorganic nutrients by the bacterial communities in our NanoSIMS measurements (Table 2).

**Table 2:** Summary of the exudate and nutrient additions to surface Eastern Mediterranean Seawater in January 2019. Values shown are the final concentration.

| Ingredient/treatment | control | control + nutrients | exudates | exudates + nutrients |
| --- | --- | --- | --- | --- |
| *Prochlorococcus* MIT9312 exudates | - | - | 25 $\mu\text{M}$ C | 25 $\mu\text{M}$ C |
| nutrients | - | 20 $\mu\text{M}$ $\text{NH}_4$,<br>2 $\mu\text{M}$ $\text{PO}_4$ | - | 20 $\mu\text{M}$ $\text{NH}_4$,<br>2 $\mu\text{M}$ $\text{PO}_4$ |
| $^{15}\text{N}$ $\text{NH}_4$ | 5 nM | 5 nM | - | - |

### Nutrient measurements

Dissolved nutrients in the cultures and for Exp. 1 were determined using a SEAL AA-3 autoanalyzer system. Nutrient concentrations from the open-ocean station Themo-2 (Exp. 2) were taken from (Reich et al., 2021) as means of the mixed period (January – April).

### Cell numbers

Bacterial cell numbers were determined by flow-cytometry. Briefly, duplicate samples were fixed with cytometry-grade glutaraldehyde (0.125% final concentration), flash-frozen with liquid nitrogen and stored at −80 °C until analyses. For measurements, samples were thawed in the dark at room temperature, stained with SYBR Green I (Molecular Probes/ ThermoFisher) for 10 min at room temperature, vortexed, and run on a BD FACSCanto™ II Flow Cytometry Analyzer Systems (BD 146 Biosciences) with 2 μm diameter fluorescent beads (Polysciences, Warminster, PA, USA) as a size and fluorescence standard. Cells were detected at Ex494nm/Em520nm (FITC channel) and by the size of cell (forward scatter). Phytoplankton cells were identified based on their cell chlorophyll (Ex482nm/Em676nm, PerCP channel) and by the size of cell (forward scatter). Flow rates were determined several times during each running session by weighing tubes with double-distilled water. Finally, the samples were analyzed with the free software “Flowing Software” (https://bioscience.fi/).

### Bacterial production (BP)

The activity of heterotrophic production was measured using the [4,5-^3^H]-leucine incorporation method (Simon et al., 1990). To this end, triplicate 1.7 ml of seawater samples were collected from each microcosm bottle, amended with 100 nmol of leucine L^−1^ (Perkin Elmer, specific activity 156 Ci mmol^−1^) and incubated for 4 h in the dark under ambient surface sea water temperature (∼19 °C). Incubations were stopped by the addition of 100 µL of ice-cold 100% trichloroacetic acid (TCA). Control samples containing seawater, radioisotope, and TCA added immediately upon incubation were also run and these reads were subtracted from the test tubes. At the lab, the samples were micro-centrifuged with TCA 5%, and 1 ml of scintillation cocktail (Ultima-Gold) was added to each vial. Disintegration per minute (DPM) were measured using a TRI-CARB 2100 TR (Packard) liquid counter. A conversion factor of 1.5 kg C mol^−1^ per mole leucine were used with an isotope dilution factor of 2 (Simon and Azam, 1989).

### Alkaline phosphatase activity (APA)

APA was determined by the 4-methylumbeliferyl phosphate (MUF-P: Sigma M8168) method according to (Thingstad and Mantoura, 2005). Substrate was added to triplicate 1 ml water samples (final concentration of 50 µM) and incubated in the dark at ambient temperature for 4 h (same as BP). The increase in fluorescence by the cleaved 4-methylumbelliferone (MUF) was measured at 365 nm excitation, 455 nm emissions (GloMax®-Multi Detection System E9032) and calibrated against a MUF standard (Sigma M1508).

### NanoSIMS analyses

Before analyses, filter pieces were covered with approximately 30 nm gold in a sputter coater (Cressington108 auto-sputter coater). SIMS imaging was performed as described in (Eigemann et al., 2019) using a NanoSIMS 50L instrument (Cameca, France). The scanning parameters were 512×512 px for areas of 20–30 μm, with a dwell time of 250 μs per pixel and a primary beam of 1 pA. All NanoSIMS measurements were analysed with the Matlab based program look@nanosims (Polerecky et al., 2012). Briefly, the 60 measured planes were checked for inconsistencies, all usable planes accumulated, regions of interest (i.e. bacterial cells) defined based on ^12^C^14^N mass pictures and ^13^C/^12^C as well as ^15^N/^14^N ratios calculated from the ion signals for each region of interest. For analyses of each measurement, first the means of background measurements were determined (i.e. regions on the filter without bacterial cells), and this mean factorized for theoretical background values (0.11 for ^13^C/^12^C and 0.00367 for ^15^N/^14^N). These factors were applied to all non-background regions of interest in the same measurement. For each treatment, measurements of different spots on the same filter as well as replicate filters (2 replicates for each treatment) were pooled.

### RNA extraction, DNA digestion, cDNA synthesis

For RNA extraction, approximately 1.5 l of each incubation bottle was filtered successively onto 5 and 0.2 µm pore-width polycarbonate filters (Exp 1), or directly onto 0.2 µm filters (Exp 2). Filters were stored in 1 ml lysis buffer (40 mM EDTA, 50 mM Tris pH 8.3, 0.75 M sucrose), flash frozen in liquid nitrogen and stored at −80 °C upon extraction. For extraction, 1 ml TRI Reagent was added to half a filter in Eppendorf tubes, vortexed, and incubated for 10 min at room temperature under rotation. Then, 200 µl chloroform were added, and filters were vortexed again and incubated for 15 min at room temperature. Following, the tubes were centrifuged at 12,000 g for 12 min at 4 °C, the supernatant transferred to fresh tubes, centrifuged again as described above, and the aqueous phase transferred to fresh tubes. Next, 1 ml ice cold 100% ethanol was added, vortexed and incubated for 1 h on ice, centrifuged at 12,000 g for 10 min, the supernatant was discarded, 1 ml ice cold 75% ethanol added, vortexed, and incubated for 10 min at −20°C. Last, tubes were centrifuged for 5 min at 7,500 g, the ethanol removed, and the RNA pellet air dried in a hood. The resulting pellet was resolved in autoclaved DEPC-treated water and quality controlled with a Nanodrop device. Remaining DNA was digested using the Turbo DNA free kit (Invitrogen) using the manufacturer’s instructions, and successful digestions tested by PCRs. RNA was transcribed into cDNA using MultiScribe reverse transcriptase following the manufacturer’s instructions (Invitrogen).

### Sequencing

Complementary DNA was PCR amplified with primers 515F and 926R (Walters et al., 2016) targeting the V4 and V5 variable regions of the microbial small subunit ribosomal RNA gene using a two-stage “targeted amplicon sequencing (TAS)” protocol (Naqib et al., 2018). Primers were modified to include linker sequences at the 5’ ends (i.e., so-called “common sequences” or CS1 and CS2 on forward and reverse primers, respectively). First stage PCR amplifications were performed in 10 microliter reactions in 96-well plates, using the MyTaq HS 2X mastermix (BioLine, Taunton, MA, USA). PCR conditions were 95°C for 5 minutes, followed by 28 cycles of 95°C for 30”, 50°C for 60” and 72°C for 90”. Subsequently, a second PCR amplification was performed in 10 microliter reactions in 96-well plates. A mastermix for the entire plate was made using the MyTaq HS 2X mastermix. Each well received a separate primer pair with a unique 10-base barcode, obtained from the Access Array Barcode Library for Illumina (Fluidigm, South San Francisco, CA; Item# 100-4876). These AccessArray primers contained the CS1 and CS2 linkers at the 3’ ends of the oligonucleotides. Cycling conditions were as follows: 95°C for 5 minutes, followed by 8 cycles of 95°C for 30”, 60°C for 30” and 72°C for 30”. A final, 7 minute elongation step was performed at 72°C. Samples were pooled in equal volume using an EpMotion5075 liquid handling robot (Eppendorf, Hamburg, Germany). The pooled library was purified using an AMPure XP cleanup protocol (0.6X, vol/vol; Agencourt, Beckmann-Coulter) to remove fragments smaller than 300 bp. The pooled libraries, with a 20% phiX spike-in, were loaded onto an Illumina MiniSeq mid-output flow cell (2×150 paired-end reads). Based on the distribution of reads per barcode, the amplicons (before purification) were re-pooled to generate a more balanced distribution of reads. The re-pooled library was purified using AMPure XP cleanup, as described above. The re-pooled libraries, with a 15% phiX spike-in, were loaded onto a MiSeq v3 flow cell, and sequenced using an Illumina MiSeq sequencer. Fluidigm sequencing primers, targeting the CS1 and CS2 linker regions, were used to initiate sequencing. Library preparation, pooling, and MiniSeq sequencing were performed at the University of Illinois at Chicago Sequencing Core (UICSQC), Research Resources Center (RRC), University of Illinois at Chicago (UIC). All forward and backward sequence reads were deposited at the European Nucleotide Archive under the accession number PRJEB44710.

### Sequence analyses

All sequences were analyzed using the Dada2 (Callahan et al., 2016) pipeline and the software packages R (R Development Team, 2020) and RStudio (RStudio Team, 2020). Briefly, forward and backward primers were trimmed, forward and backward reads truncated after quality inspections to 280 and 210 bases, respectively, and after merging of forward and backward sequences, a consensus length only between 404 and 417 bases was accepted. For taxonomic assignment, Silva database version 138 (Quast et al., 2013) was used. All chloroplasts, mitochondria, archaea, eukaryotes and Amplicon sequence variants (ASVs) with no found relatives were discarded from downstream analyses. The complete ASV table with absolute read counts, sequences, sequence lengths as well as meta data for all samples is accessible as Supplemental 9. After inspections of rarefaction curves, all samples with less than 2509 reads were discarded from further analyses, and all remaining samples subsampled to 2509 reads.

### Statistical analyses

Bacterial communities were analyzed with Shannon diversities, Non-metric-multidimensional-scaling (NMDS), and heatmaps of the most abundant ASVs. NMDS was performed using the ‘metaMDS’ command and the bray distance in the ‘vegan’ package (Oksanen et al. 2019) in order to analyze differences between the communities based on the subsampled absolute ASV tables. An analysis of similarities (ANOSIM), which tests for significant differences between communities, was conducted using the ‘vegan’ package. Heatmaps for the 40 most abundant ASVs (subsampled and separately for both experiments) were calculated using the heatmap 3 package (Zhao et al., 2014). Homogeneities of variances were tested by Bartlett or Levene’s (Shannon diversity) tests. Differences between the different treatments in terms of Shannon diversity, incorporation of ^13^C and ^15^N, cell numbers, and bacterial production were tested by ANOVAs with subsequent Tukey post-hoc tests if homogeneity of variances was given. If homogeneity was not given, Kruskal-Wallis tests were calculated, with subsequent Tukey-Nemenyi post-hoc tests. To test for specific ASVs associated with treatments, LDA effect size (LEfSe) analyses (Segata et al., 2011) were executed with the online tool https://huttenhower.sph.harvard.edu/galaxy. For the open-ocean community, treatments were assigned as class without subclass, whereas for the open-ocean community, treatments were assigned as class, and the fraction (free and attached) assigned as subclass. For this multi-class analysis, a “one-against-all” strategy was applied. All analyses, except denoted other, were performed with R (R Development Team, 2020) and RStudio (RStudio Team, 2020), all graphics were executed with the ggplot2 package (Ginestet, 2011), and refined with the freeware Inkscape (https://inkscape.org).

## Acknowledgement

We thank the captain and crew of the R/V Mediterranean Explorer (EcoOcean) for help at sea, Mike Krom, Anat Tsemel and Tal Ben-Ezra for the inorganic nutrient analysis, and Stefan Green (DNA Services Facility at the University of Illinois at Chicago) for the amplicon sequencing. We also thank Tom Reich, Dalit Roth-Rosenberg, Tal Luzzatto-Knaan, Noam Nago, and Natalia Belkin for excellent help with the experiments, and Annett Grüttmüller for NanoSIMS measurements. This work was supported by the Human Frontier Science Program (HFSP) through the grant number RGB 0020/2016 (DS, MV and HPG), by the National Science Foundation - United States-Israel Binational Science Foundation Program in Oceanography (grant number 1635070/2016532 to DS) and by the Israel Ministry of Science and Technology (grant number 3-17404 to DS). The experiment at THEMO-2 was performed as part of the SoMMoS (Southeastern Mediterranean Monthly cruise Series) project, with ship-time funded by the Leon H. Charney School of Marine Sciences with help from EcoOcean and IOLR. The NanoSIMS at the Leibnitz-Institute for Baltic Sea research in Warnemuende (IOW) was funded by the German Federal Ministry of Education and Research (BMBF), grant identifier 03F0626A.

## References

Amin, S.A., Parker, M.S., and Armbrust, E.V. (2012) Interactions between diatoms and bacteria. Microbiol Mol Biol Rev 76: 667–684.

Amin, S.A., Hmelo, L.R., van Tol, H.M., Durham, B.P., Carlson, L.T., Heal, K.R. et al. (2015) Interaction and signalling between a cosmopolitan phytoplankton and associated bacteria. Nature 522: 98–101.

Azam, F., and Malfatti, F. (2007) Microbial structuring of marine ecosystems. Nat Rev Microbiol 5: 782–791.

Beier, S., Rivers, A.R., Moran, M.A., and Obernosterer, I. (2015) The transcriptional response of prokaryotes to phytoplankton-derived dissolved organic matter in seawater. Environ Microbiol 17: 3466–3480.

Beliaev, A.S., Romine, M.F., Serres, M., Bernstein, H.C., Linggi, B.E., Markillie, L.M. et al. (2014) Inference of interactions in cyanobacterial-heterotrophic co-cultures via transcriptome sequencing. ISME J 8: 2243–2255.

Berman, T., and Bronk, D.A. (2003) Dissolved organic nitrogen: a dynamic participant in aquatic ecosystems. Aquatic Microbial Ecology 31: 279–305.

Bertilsson, S., Berglund, O., Pullin, M., and Chisholm, S. (2005) Release of dissolved organic matter by Prochlorococcus. Vie Et Milieu-life and Environment 55: 225–231.

Bertocchi, C., Navarini, L., Cesàro, A., and Anastasio, M. (1990) Polysaccharides from cyanobacteria. Carbohydrate Polymers 12: 127–153.

Bhatnagar, M., and Bhatnagar, A. (2019) Diversity of Polysaccharides in Cyanobacteria. In Microbial Diversity in Ecosystem Sustainability and Biotechnological Applications: Volume 1 Microbial Diversity in Normal & Extreme Environments. Satyanarayana, T., Johri, B.N., and Das, S.K. (eds). Singapore: Springer Singapore, pp. 447–496.

Bianchi, A., Van Wambeke, F., and Garcin, J. (1998) Bacterial utilization of glucose in the water column from eutrophic to oligotrophic pelagic areas in the eastern North Atlantic Ocean. Journal of Marine Systems 14: 45–55.

Biller, S.J., Berube, P.M., Lindell, D., and Chisholm, S.W. (2015) Prochlorococcus: the structure and function of collective diversity. Nature Reviews Microbiology 13: 13–27.

Bradley, P.B., Sanderson, M.P., Nejstgaard, J.C., Sazhin, A.F., Frischer, M.E., Killberg-Thoreson, L.M. et al. (2010) Nitrogen uptake by phytoplankton and bacteria during an induced Phaeocystis pouchetii bloom, measured using size fractionation and flow cytometric sorting. Aquatic Microbial Ecology 61: 89–104.

Buchan, A., LeCleir, G.R., Gulvik, C.A., and Gonzalez, J.M. (2014) Master recyclers: features and functions of bacteria associated with phytoplankton blooms. Nat Rev Microbiol 12: 686–698.

Callahan, B.J., McMurdie, P.J., Rosen, M.J., Han, A.W., Johnson, A.J.A., and Holmes, S.P. (2016) DADA2: High-resolution sample inference from Illumina amplicon data. Nature methods 13: 581–583.

Carlson, C.A., and Ducklow, H.W. (1996) Growth of bacterioplankton and consumption of dissolved organic carbon in the Sargasso Sea. Aquatic Microbial Ecology 10: 69–85.

Carlson, C.A., Giovannoni, S.J., Hansell, D.A., Goldberg, S.J., Parsons, R., and Vergin, K. (2004) Interactions among dissolved organic carbon, microbial processes, and community structure in the mesopelagic zone of the northwestern Sargasso Sea. Limnology and Oceanography 49: 1073–1083.

Christie-Oleza, J.A., Sousoni, D., Lloyd, M., Armengaud, J., and Scanlan, D.J. (2017) Nutrient recycling facilitates long-term stability of marine microbial phototroph-heterotroph interactions. Nat Microbiol 2: 17100.

Cole, J.J. (1982) Interactions between bacteria and algae in aquatic ecosystems. Annual Review of Ecology and Systematics 13: 291–314.

Eigemann, F., Vogts, A., Voss, M., Zoccarato, L., and Schulz-Vogt, H. (2019) Distinctive tasks of different cyanobacteria and associated bacteria in carbon as well as nitrogen fixation and cycling in a late stage Baltic Sea bloom. PLoS One 14: e0223294.

Eilers, H., Pernthaler, J., Glöckner, F.O., and Amann, R. (2000) Culturability and in situ abundance of pelagic bacteria from the North Sea. Applied and Environmental Microbiology 66: 3044–3051.

Field, C.B., Behrenfeld, M.J., Randerson, J.T., and Falkowski, P. (1998) Primary production of the biosphere: Integrating terrestrial and oceanic components. Science 281: 237–240.

Flombaum, P., Gallegos, J.L., Gordillo, R.A., Rincón, J., Zabala, L.L., Jiao, N. et al. (2013) Present and future global distributions of the marine Cyanobacteria Prochlorococcus and Synechococcus. Proceedings of the National Academy of Sciences 110: 9824–9829.

Fouilland, E., Tolosa, I., Bonnet, D., Bouvier, C., Bouvier, T., Bouvy, M. et al. (2014) Bacterial carbon dependence on freshly produced phytoplankton exudates under different nutrient availability and grazing pressure conditions in coastal marine waters. FEMS Microbiol Ecol 87: 757–769.

Ginestet, C. (2011) ggplot2: Elegant Graphics for Data Analysis. Journal of the Royal Statistical Society: Series A (Statistics in Society) 174: 245–246.

Gobet, A., Barbeyron, T., Matard-Mann, M., Magdelenat, G., Vallenet, D., Duchaud, E., and Michel, G. (2018) Evolutionary evidence of algal polysaccharide degradation acquisition by Pseudoalteromonas carrageenovora 9(T) to adapt to macroalgal niches. Front Microbiol 9: 2740.

Grossart, H.-P. (1999) Interactions between marine bacteria and axenic diatoms (Cylindrotheca fusiformis, Nitzschia laevis, and Thalassiosira weissflogii) incubated under various conditions in the lab. Aquatic Microbial Ecology 19: 1–11.

Grossart, H.-P., and Simon, M. (2007) Interactions of planktonic algae and bacteria: effects on algal growth and organic matter dynamics. Aquatic Microbial Ecology 47: 163–176.

Grossart, H.P., Tang, K.W., Kiorboe, T., and Ploug, H. (2007) Comparison of cell-specific activity between free-living and attached bacteria using isolates and natural assemblages. FEMS Microbiol Lett 266: 194–200.

Grossowicz, M., Roth-Rosenberg, D., Aharonovich, D., Silverman, J., Follows, M.J., and Sher, D. (2017) Prochlorococcus in the lab and in silico: The importance of representing exudation. Limnology and Oceanography 62: 818–835.

Grzymski, J.J., and Dussaq, A.M. (2012) The significance of nitrogen cost minimization in proteomes of marine microorganisms. The ISME Journal 6: 71–80.

Hazan, O., Silverman, J., Sisma-Ventura, G., Ozer, T., Gertman, I., Shoham-Frider, E. et al. (2018) Mesopelagic prokaryotes alter surface phytoplankton production during simulated deep mixing experiments in Eastern Mediterranean Sea waters. Frontiers in Marine Science 5.

Huang, Y., Nicholson, D., Huang, B., and Cassar, N. (2021) Global estimates of marine gross primary production based on machine learning upscaling of field observations. Global Biogeochemical Cycles 35: e2020GB006718.

Johnson, Z.I., Zinser, E.R., Coe, A., McNulty, N.P., Woodward, E.M.S., and Chisholm, S.W. (2006) Niche partitioning among Prochlorococcus ecotypes along ocean-scale environmental gradients. Science 311: 1737–1740.

Karlson, A.M., Duberg, J., Motwani, N.H., Hogfors, H., Klawonn, I., Ploug, H. et al. (2015) Nitrogen fixation by cyanobacteria stimulates production in Baltic food webs. Ambio 44 Suppl 3: 413–426.

Klippel, B., Lochner, A., Bruce, D.C., Davenport, K.W., Detter, C., Goodwin, L.A. et al. (2011) Complete genome sequence of the marine cellulose- and xylan-degrading bacterium Glaciecola sp. strain 4H-3-7+YE-5. J Bacteriol 193: 4547–4548.

Krom, M., Kress, N., Berman-Frank, I., and Rahav, E. (2014) Past, present and future patterns in the nutrient chemistry of the Eastern Mediterranean. In The Mediterranean Sea: Its history and present challenges. Goffredo, S., and Dubinsky, Z. (eds). Dordrecht: Springer Netherlands, pp. 49–68.

Krom, M.D., Emeis, K.C., and Van Cappellen, P. (2010) Why is the Eastern Mediterranean phosphorus limited? Progress in Oceanography 85: 236–244.

Lidbury, I.D., Murrell, J.C., and Chen, Y. (2015) Trimethylamine and trimethylamine N-oxide are supplementary energy sources for a marine heterotrophic bacterium: implications for marine carbon and nitrogen cycling. The ISME Journal 9: 760–769.

Liefer, J.D., Garg, A., Fyfe, M.H., Irwin, A.J., Benner, I., Brown, C.M. et al. (2019) The Macromolecular Basis of Phytoplankton C:N:P Under Nitrogen Starvation. Front Microbiol 10: 763.

Livanou, E., Lagaria, A., Psarra, S., and Lika, K. (2017) Dissolved organic matter release by phytoplankton in the context of the Dynamic Energy Budget theory. Biogeosciences 42: 1–33.

Lombard, V., Golaconda Ramulu, H., Drula, E., Coutinho, P.M., and Henrissat, B. (2014) The carbohydrate-active enzymes database (CAZy) in 2013. Nucleic Acids Res 42: D490–495.

McCarren, J., Becker, J.W., Repeta, D.J., Shi, Y., Young, C.R., Malmstrom, R.R. et al. (2010) Microbial community transcriptomes reveal microbes and metabolic pathways associated with dissolved organic matter turnover in the sea. PNAS 107: 16420–16427.

Mella-Flores, D., Mazard, S., Humily, F., Partensky, F., Mahé, F., Bariat, L. et al. (2011) Is the distribution of Prochlorococcus and Synechococcus ecotypes in the Mediterranean Sea affected by global warming? Biogeosciences 8: 2785–2804.

Meon, B., and Kirchman, D.L. (2001) Dynamics and molecular composition of dissolved organic material during experimental phytoplankton blooms. Marine Chemistry 75: 185–199.

Moore, C.M., Mills, M.M., Arrigo, K.R., Berman-Frank, I., Bopp, L., Boyd, P.W. et al. (2013) Processes and patterns of oceanic nutrient limitation. Nature Geoscience 6: 701–710.

Morán, X.A., G., Josep, M.G., Pedros-Alio, C., and Marta, E. (2002) Partitioning of phytoplanktonic organic carbon production and bacterial production along a coastal-offshore gradient in the NE Atlantic during different hydrographic regimes. Aquatic Microbial Ecology 29: 239–252.

Mühlenbruch, M., Grossart, H.P., Eigemann, F., and Voss, M. (2018) Mini-review: Phytoplankton-derived polysaccharides in the marine environment and their interactions with heterotrophic bacteria. Environ Microbiol 20: 2671–2685.

Naqib, A., Poggi, S., Wang, W., Hyde, M., Kunstman, K., and Green, S.J. (2018) Making and Sequencing Heavily Multiplexed, High-Throughput 16S Ribosomal RNA Gene Amplicon Libraries Using a Flexible, Two-Stage PCR Protocol. Methods Mol Biol 1783: 149–169.

Nemergut, D.R., Schmidt, S.K., Fukami, T., O’Neill, S.P., Bilinski, T.M., Stanish, L.F. et al. (2013) Patterns and processes of microbial community assembly. Microbiology and Molecular Biology Reviews 77: 342–356.

Ofaim, S., Sulheim, S., Almaas, E., Sher, D., and Segrè, D. (2021) Dynamic allocation of carbon storage and nutrient-dependent exudation in a revised genome-scale model of Prochlorococcus. Frontiers in Genetics 12.

Partensky, F., Hess, W.R., and Vaulot, D. (1999) Prochlorococcus, a marine photosynthetic prokaryote of global significance. Microbiology and molecular biology reviews : MMBR 63: 106–127.

Pedler, B.E., Aluwihare, L.I., and Azam, F. (2014) Single bacterial strain capable of significant contribution to carbon cycling in the surface ocean. Proceedings of the National Academy of Sciences 111: 7202–7207.

Pheng, S., Ayyadurai, N., Park, A.Y., and Kim, S.G. (2017) Psychrosphaera aquimarina sp. nov., a marine bacterium isolated from seawater collected from Asan Bay, Republic of Korea. Int J Syst Evol Microbiol 67: 4820–4824.

Pinhassi, J., Laura, G.-C., Laura, A.-S., Maria, M.S., Montserrat, V., Carlos, P.-A., and Josep, M.G. (2006) Seasonal changes in bacterioplankton nutrient limitation and their effects on bacterial community composition in the NW Mediterranean Sea. Aquatic Microbial Ecology 44: 241–252.

Piontek, J., Handel, N., De Bodt, C., Harlay, J., Chou, L., and Engel, A. (2011) The utilization of polysaccharides by heterotrophic bacterioplankton in the Bay of Biscay (North Atlantic Ocean). Journal of Plankton Research 33: 1719–1735.

Polerecky, L., Adam, B., Milucka, J., Musat, N., Vagner, T., and Kuypers, M.M. (2012) Look@NanoSIMS--a tool for the analysis of nanoSIMS data in environmental microbiology. Environ Microbiol 14: 1009–1023.

Quast, C., Pruesse, E., Yilmaz, P., Gerken, J., Schweer, T., Yarza, P. et al. (2013) The SILVA ribosomal RNA gene database project: improved data processing and web-based tools. Nucleic acids research 41: D590–D596.

R Development Team (2020) R: A language and environment for statistical computing. R Foundation for Statistical Computing, Vienna, Austria. URL https://www.R-project.org/. In: R Foundation for Statistical Computing.

Rahav, E., Silverman, J., Raveh, O., Hazan, O., Rubin-Blum, M., Zeri, C. et al. (2019) The deep water of Eastern Mediterranean Sea is a hotspot for bacterial activity. Deep Sea Research Part II: Topical Studies in Oceanography 164: 135–143.

Raveh, O., David, N., Rilov, G., and Rahav, E. (2015) The temporal dynamics of coastal phytoplankton and bacterioplankton in the Eastern Mediterranean Sea. PLOS ONE 10.

Reich, T., Ben-Ezra, T., Belkin, N., Tsemel, A., Aharonovich, D., Roth-Rosenberg, D. et al. (2021) Seasonal dynamics of phytoplankton and bacterioplankton at the ultra-oligotrophic southeastern Mediterranean Sea. bioRxiv: 2021.2003.2024.436734.

Ribalet, F., Swalwell, J., Clayton, S., Jiménez, V., Sudek, S., Lin, Y. et al. (2015) Light-driven synchrony of Prochlorococcus growth and mortality in the subtropical Pacific gyre. Proceedings of the National Academy of Sciences 112: 8008–8012.

Riemann, L., Holmfeldt, K., and Titelman, J. (2009) Importance of viral lysis and dissolved DNA for bacterioplankton activity in a P-limited estuary, Northern Baltic Sea. Microb Ecol 57: 286–294.

Romera-Castillo, C., Sarmento, H., Alvarez-Salgado, X.A., Gasol, J.M., and Marrasé, C. (2011) Net production and consumption of fluorescent colored dissolved organic matter by natural bacterial assemblages growing on marine phytoplankton exudates. Applied and Environmental Microbiology 77: 7490–7498.

Rooney-Varga, J.N., Giewat, M.W., Savin, M.C., Sood, S., LeGresley, M., and Martin, J.L. (2005) Links between phytoplankton and bacterial community dynamics in a coastal marine environment. Microb Ecol 49: 163–175.

Roth-Rosenberg, D., Aharonovich, D., Omta, A.W., Follows, M.J., and Sher, D. (2021a) Dynamic macromolecular composition and high exudation rates in Prochlorococcus. Limnology and Oceanography 66: 1759–1773.

Roth-Rosenberg, D., Haber, M., Goldford, J., Lalzar, M., Aharonovich, D., Al-Ashhab, A. et al. (2021b) Particle-associated and free-living bacterial communities in an oligotrophic sea are affected by different environmental factors. Environmental Microbiology n/a.

RStudio Team (2020) RStudio: Integrated Development for R. RStudio, PBC, Boston, MA. URL https://www.rstudio.com/.

Saito, M.A., McIlvin, M.R., Moran, D.M., Goepfert, T.J., DiTullio, G.R., Post, A.F., and Lamborg, C.H. (2014) Multiple nutrient stresses at intersecting Pacific Ocean biomes detected by protein biomarkers. Science 345: 1173–1177.

Sarmento, H., and Gasol, J.M. (2012) Use of phytoplankton-derived dissolved organic carbon by different types of bacterioplankton. Environ Microbiol 14: 2348–2360.

Sarmento, H., Morana, C., and Gasol, J.M. (2016) Bacterioplankton niche partitioning in the use of phytoplankton-derived dissolved organic carbon: quantity is more important than quality. ISME J 10: 2582–2592.

Sarmento, H., Romera-Castillo, C., Lindh, M., Pinhassi, J., Sala, M.M., Gasol, J.M. et al. (2013) Phytoplankton species-specific release of dissolved free amino acids and their selective consumption by bacteria. Limnology and Oceanography 58: 1123–1135.

Segata, N., Izard, J., Waldron, L., Gevers, D., Miropolsky, L., Garrett, W.S., and Huttenhower, C. (2011) Metagenomic biomarker discovery and explanation. Genome Biol 12: R60.

Seymour, J.R., Ahmed, T., and Stocker, R. (2009) Bacterial chemotaxis towards the extracellular products of the toxic phytoplankton Heterosigma akashiwo. Journal of Plankton Research 31: 1557–1561.

Seymour, J.R., Ahmed, T., Durham, W.M., and Stocker, R. (2010) Chemotactic response of marine bacteria to the extracellular products of Synechococcus and Prochlorococcus. Aquatic Microbial Ecology 59: 161–168.

Seymour, J.R., Amin, S.A., Raina, J.B., and Stocker, R. (2017) Zooming in on the phycosphere: the ecological interface for phytoplankton-bacteria relationships. Nat Microbiol 2: 17065.

Sharma, A.K., Becker, J.W., Ottesen, E.A., Bryant, J.A., Duhamel, S., Karl, D.M. et al. (2014) Distinct dissolved organic matter sources induce rapid transcriptional responses in coexisting populations of Prochlorococcus, Pelagibacter and the OM60 clade. Environ Microbiol 16: 2815–2830.

Shaw, S.L., Chisholm, S.W., and Prinn, R.G. (2003) Isoprene production by Prochlorococcus, a marine cyanobacterium, and other phytoplankton. Marine Chemistry 80: 227–245.

Simon, M., and Azam, F. (1989) Protein content and protein synthesis rates of planktonic marine bacteria. Marine Ecology Progress Series 51: 201–213.

Simon, M., Alldredge, A.L., and Azam, F. (1990) Bacterial carbon dynamics on marine snow. Marine Ecology Progress Series 65: 205–211.

Sisma-Ventura, G., and Rahav, E. (2019) DOP stimulates heterotrophic bacterial production in the oligotrophic southeastern Mediterranean coastal waters. Frontiers in Microbiology 10.

Skoog, A., Whitehead, K., Sperling, F., and Junge, K. (2002) Microbial glucose uptake and growth along a horizontal nutrient gradient in the North Pacific. Limnology and Oceanography 47: 1676–1683.

Taylor, J.D., and Cunliffe, M. (2017) Coastal bacterioplankton community response to diatom-derived polysaccharide microgels. Environ Microbiol Rep 9: 151–157.

Teeling, H., Fuchs, B.M., Bennke, C.M., Kruger, K., Chafee, M., Kappelmann, L. et al. (2016) Recurring patterns in bacterioplankton dynamics during coastal spring algae blooms. Elife 5: e11888.

Teeling, H., Fuchs, B.M., Becher, D., Klockow, C., Gardebrecht, A., Bennke, C.M. et al. (2012) Substrate-controlled succession of marine bacterioplankton populations induced by a phytoplankton bloom. Science 336: 608–611.

Thingstad, T.F., and Mantoura, R.F.C. (2005) Titrating excess nitrogen content of phosphorous-deficient eastern Mediterranean surface water using alkaline phosphatase activity as a bio-indicator. Limnology and Oceanography Methods 3: 94–100.

Thornton, D.C.O. (2014) Dissolved organic matter (DOM) release by phytoplankton in the contemporary and future ocean. European Journal of Phycology 49: 20–46.

Walters, W., Hyde, E.R., Berg-Lyons, D., Ackermann, G., Humphrey, G., Parada, A. et al. (2016) Improved bacterial 16S rRNA gene (V4 and V4-5) and fungal internal transcribed spacer marker gene primers for microbial community surveys. mSystems 1.

Wu, X., Wang, Y., Zhu, Y., Tian, H., Qin, X., Cui, C. et al. (2019) Variability in the response of bacterial community assembly to environmental selection and biotic factors depends on the immigrated bacteria, as revealed by a soil microcosm experiment. mSystems 4: e00496–00419.

York, A. (2018) Marine biogeochemical cycles in a changing world. Nature Reviews Microbiology 16: 259–259.

Zhao, S., Guo, Y., Sheng, Q., and Shyr, Y. (2014) Heatmap3: an improved heatmap package with more powerful and convenient features. BMC Bioinformatics 15: P16.

