## Supplementary figures and images for "Phytoplankton exudates provide full nutrition to a subset of accompanying heterotrophic bacteria via carbon, nitrogen and phosphorus allocation"

### Supplemental 5

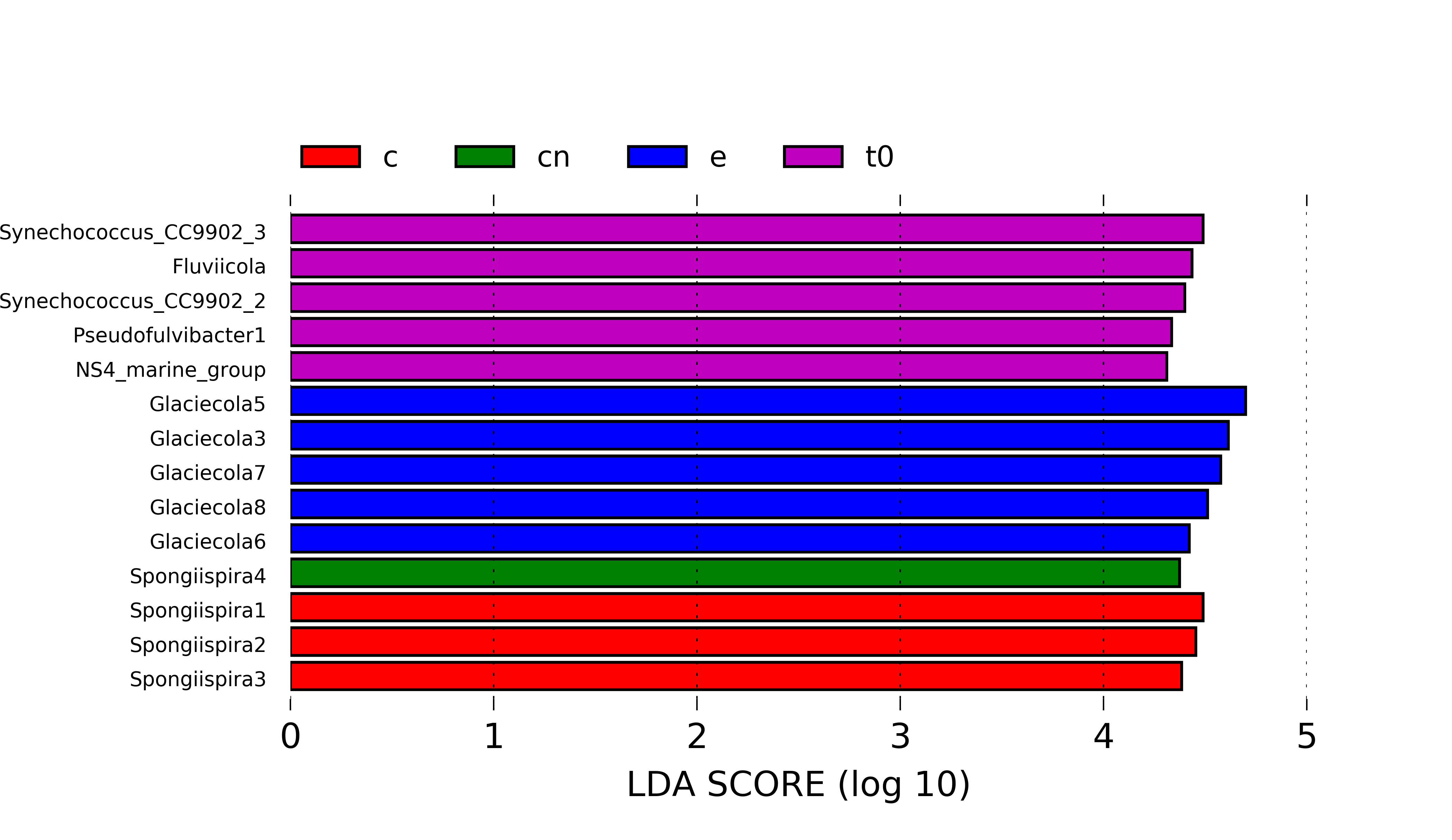

### Supplemental 7

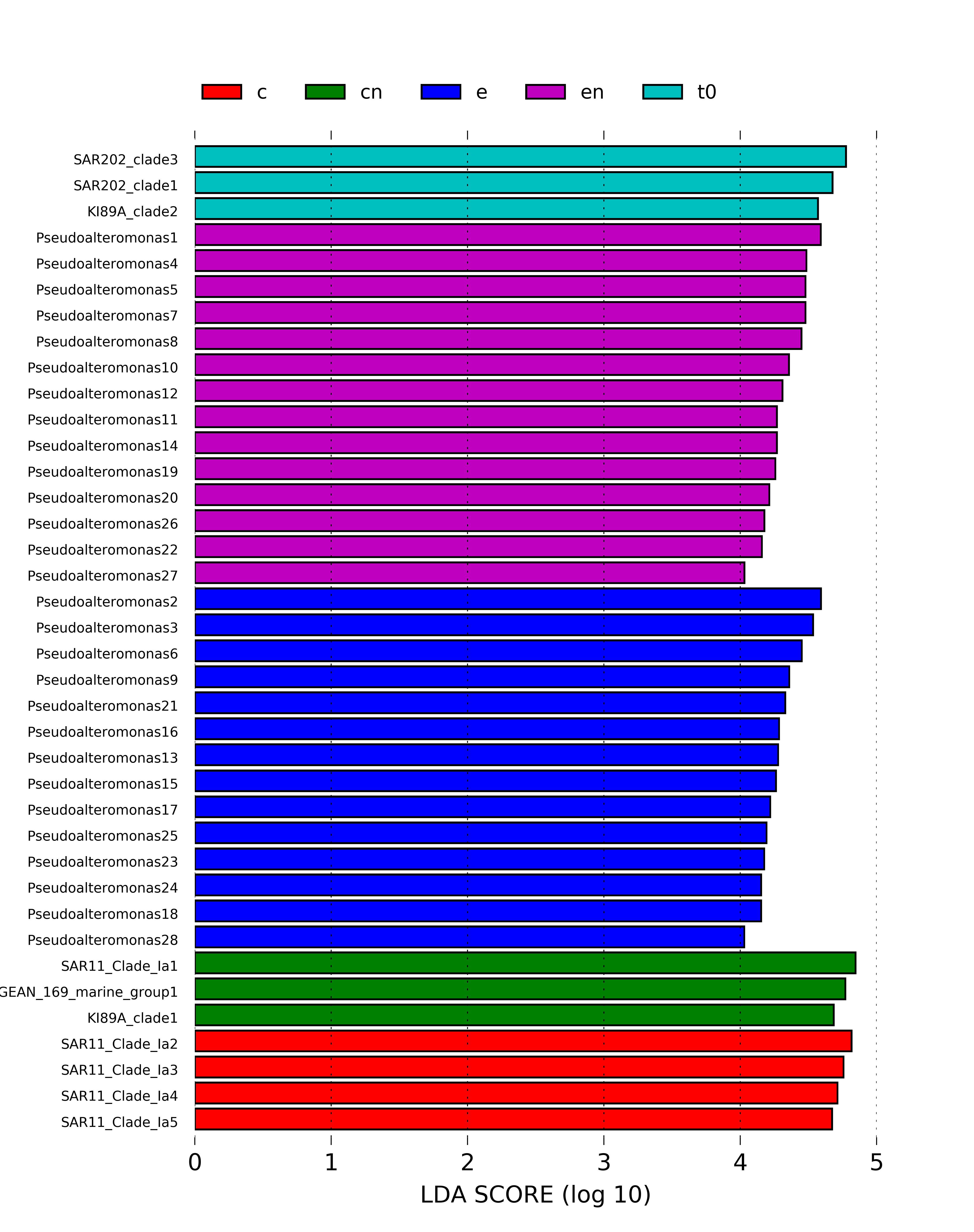
